# An open-source experimental framework for automation of high-throughput cell biology experiments

**DOI:** 10.1101/2020.07.02.185454

**Authors:** Pavel Katunin, Ashley Cadby, Anton Nikolaev

## Abstract

Modern data analysis methods, such as optimisation algorithms or machine and deep learning, have been successfully applied to a number of biological, biotechnological and medical questions. For these methods to be efficient, a large number of high quality experiments need to be conducted, which requires a high degree of automation. Here we report an open-source hardware that allows for automatic high-throughput generation of large amounts of cell biology data. The hardware consists of an automatic XY-stage for moving a multiwell plate containing growing cells; a perfusion manifold allowing application of up to 8 different solutions; and a small epifluorescent microscope. It is extremely cheap (approximately £400 without and £2500 with a fluorescent microscope) and is easily customizable for individual experimental needs. We demonstrate the usability of this platform with high-throughput Ca2+ imaging and large-scale labelling experiments.

**Key points:**

- We present an open source framework for automation of cell biology experiments
- The framework consists of an XY platform, application of up to 8 solutions and a small epifluorescent microscope with autofocusing
- Very cheap (£400 without a fluorescent microscope and £2500 with a fluorescent microscope), customisable,
- Can be used in a variety of biological applications such as imaging of fluorescent reporters, optimisation of treatment conditions and fluorescent labelling

## INTRODUCTION

Deep learning and artificial neural networks (ANN) developed in the past decade have been proven useful for image analysis, optimisation tasks and robotics[1-3]. They are also becoming increasingly popular in solving biological problems. For example, ANN based algorithms of cell segmentation are more accurate and much faster than conventional methods[4]. Deep learning also helps to detect transformed cells in human tissues[5, 6], optimise treatment conditions[7] and explain animal behaviour[8]. Recently, an online platform has been developed to allow researchers without any prior knowledge of deep learning to use it in their own applications, further increasing the usefulness of deep learning as an analytical tool [9].

One important consideration that has to be taken into account when applying deep learning and other machine learning methods is the size of the training datasets. Typically, deep learning requires thousands or tens of thousands of data points[10]. This is often not feasible in biological experiments as they often take a long time to conduct. Resultantly, there is a demand for automated systems that would allow hundreds or thousands of experiments to be performed simultaneously. Commercially available automation tools are useful, however these are expensive (>$10000-15000), large, non-flexible and mainly focus on solution handling. Importantly, experimental automation requires image analysis and application of solutions performed simultaneously, while many commercially available microscopes provide limited possibility for such closed loop experiments. Therefore, there is a demand for an open-source system that combines high-throughput microscopy in multiwell plates, automated solution application and simultaneous data analysis.

Here we developed an open-source experimental framework allowing for automated cell biology experiments. It is cheap, easily customisable and allows for up to 384 experiments to be performed simultaneously or sequentially. Its small epifluorescent microscope can be used to perform imaging of fluorescent reporters (e.g. GCaMP and synthetic calcium dyes, [11]) or samples labelled with fluorescent antibodies or dyes. The proposed experimental framework will make generation of biological data faster, more accurate and on a larger scale.

## RESULTS

### 1. Platform description

#### 1.1 Hardware

The hardware (Fig.1A) consists of several principal modules: the X-Y stage (Fig.1B) moving a multiwell plate horizontally; a small epifluorescent microscope (Fig 1C) with autofocusing system (Fig. 1D) and a perfusion manifold performing application of 8 solutions into individual wells (Fig.1E, F). Building instructions and all files for 3D printing are available at https://github.com/frescolabs/FrescoM (see also Supplementary Text).

**Fig 1.**
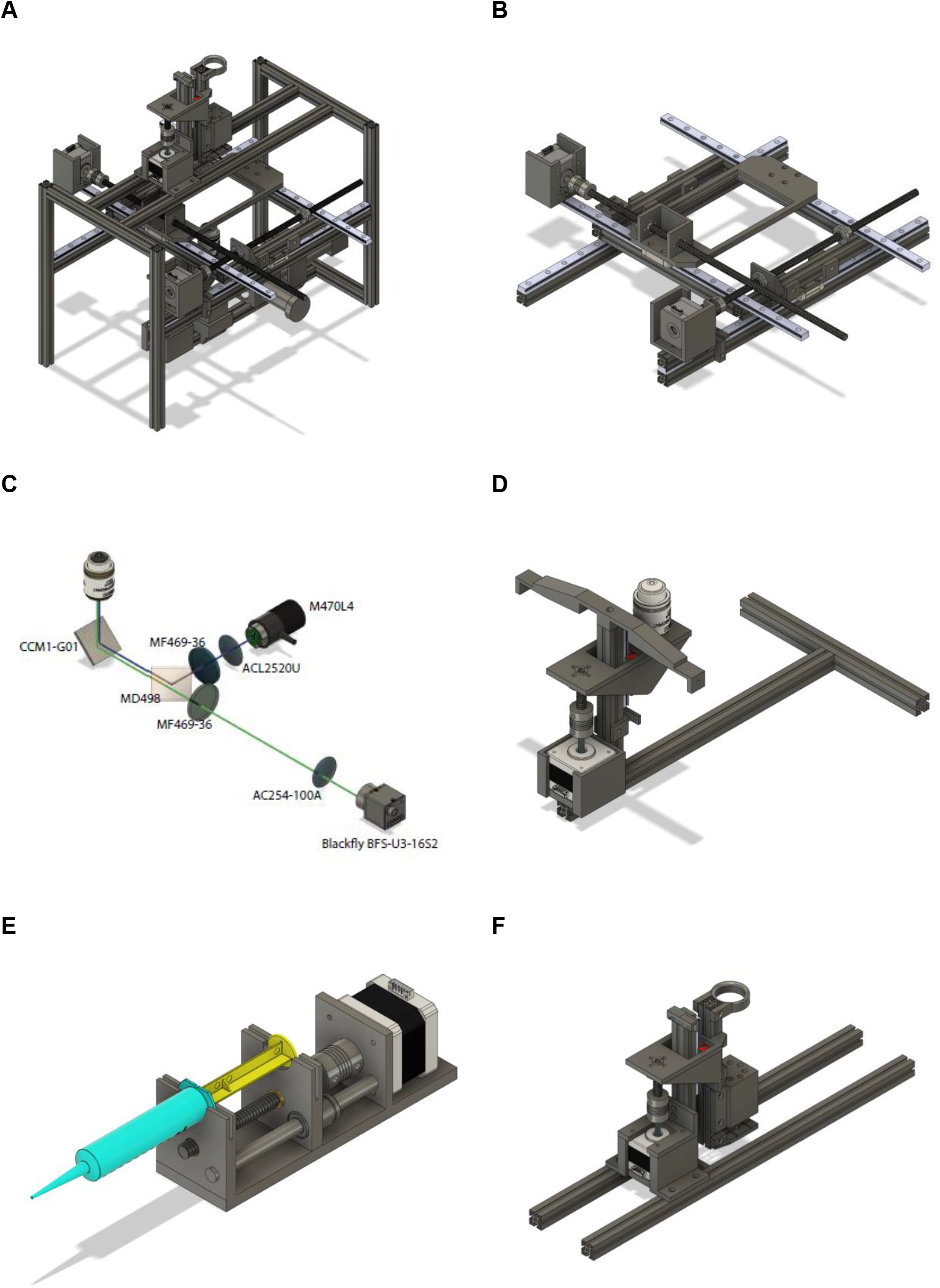
Overview of the hardware design. A,-Assembled hardware consists of 3 modules: XY platform moving in horizontal directions; perfusion manifold moving up and down, epifluorescent microscope with a Z-stage. B - The XY-stage consists of 2 side frames made of aluminium extrusions. Two long horizontal extrusions hold MGH12 rails forming the X-axis. A Nema 17 motor is attached to the side frame and moves the plate along the X axis. Rotation of the stepper motor moves the leading screw that is attached to either the top platform or to a plate holder. The Y stage is based on an aluminium frame holding the plate holder. The Y-axis motor is attached directly to one of the rails. C, D-Optical design of the microscope and the autofocusing system. Numbers represent Thorlabs (all optics) and Flir (camera) item numbers. The system implements a standard inverted fluorescent microscope with the objective attached separately to a vertically oriented linear actuator driven by a Nema 17 motor (D). E-design of a syringe pump. The main platform holds one Nema 17 motor, lead screw and an 8 mm rod. The moving part represents a holder for a nut and linear bearing. F-perfusion manifold for 8 syringe pumps. Manifold holds 8 pipette tips with one connected to the peristaltic pump. The latter is located slightly higher than others, which helps to keep the volume in each well constant. The manifold is fixed on a linear actuator and is driven by a Nema 17 motor. The whole system is fixed on two long horizontal extrusions attached to the top of the main frame. Four white LEDs connected in series are attached to the bottom of the manifold to provide bright field microscope functionality.

The overall structure is built with MakerbeamXL 15×15mm extrusions connected with each other either by L- and T-shaped aluminium brackets or 3D printed parts. We found that using alluminium extrusion frame instead of fully 3D printed parts (both models available on https://github.com/frescolabs/FrescoM) makes the whole system more stable and reduces the vibration (data not shown). The frame consists of 8 side extrusions (4 vertical 300mm and 3 horizontal 200mm, 2 at the top and 1 at the bottom, Fig 1A). The sides are connected with two 400mm extrusions holding MGH12 rails driving the X-axis. The Y-axis (Fig. 1B) resides on a square frame constructed of four 200mm extrusions and connected to the X-axis by 2 3D printed holders. Two MGH12 rails are positioned on the frame and hold one Nema17 motor and a 96 well plate holder (Fig. 1B).

The perfusion manifold (Fig.1F) consists of a holder for 8 gel loading tips attached to flexible tubings via luer connectors. One of the tips is connected to a peristaltic pump. It is located higher than the application tips, thus providing constant solution height in each well. Alternatively, the peristaltic pump tip can be placed lower than others thus providing more efficient washout of solution with a smaller volume (see section 2, and Fig.3B). Solution change in individual wells is achieved by perfusing the wells with three to five well volumes or removal of the old solution and subsequent addition of a new solution. The perfusion manifold is connected to a vertical 150mm extrusion, which is connected to a vertically oriented linear actuator consisting of MGN12 rail, T8 lead screw and a Nema 17 motor. The actuator is attached to two horizontal extrusions. Standard syringe pumps (Fig. 1E) are used for solution application. The manifold is designed to have a modular structure – other modules can be attached below, above or instead of the perfusion module. For example, we have designed a set of 4 white LEDs to be attached at the bottom of the manifold for bright field microscopy as well as a holding ring for additional tubing attached at the top. Other modules, such as an electrode holder for simple electrophysiological experiments; holder for a miniature light guide for spatially controlled optogenetics experiments; or minipumps for individual cell manipulation can be designed and integrated for additional experimental customisation.

The schematics of the fluorescent microscope are shown in Figs 1C and D. It is a standard inverted fluorescent microscopy system with the following key features. We use a non-infinity corrected objective (or, alternatively, small aspheric lens, f=2.75, NA=0.64) and a 100 mm camera lens (D=25.4 mm). The GFP cube consists of blue and green Thorlabs filters (MF469-35 and MF525-39) and a dichroic mirror (MD498, Thorlabs). The objective is not attached to the rest of the microscope but is moving separately in a Z-stage (Fig. 1D), connected to the main frame via an aluminium extrusion. If the size of the sample is an issue, a more expensive infinity corrected objective can be used instead. In this case, there will be no fluctuations in the estimated size when focusing varies from well to well.

#### 1.2 Automatic control and electric circuits

All motors are operated via 3 CNC shields connected to Arduino Mega via a PCB board (Fig. 2A and Supplementary Figure 3). Shield 1 operates the XY stage, perfusion module and autofocusing (Arduino pins 5-12 and enabling pin 13). Shields 2 and 3 operate 8 perfusion pumps (Arduino pins 23-53, odd numbers). In order for the software to have accurate estimates of the position of each axis and pump, endstops are attached to the rail of each axis (Arduino pins 22-44, even numbers). The Gerber file for PCB generation can be downloaded from https://github.com/frescolabs/FrescoM.

**Fig 2.**
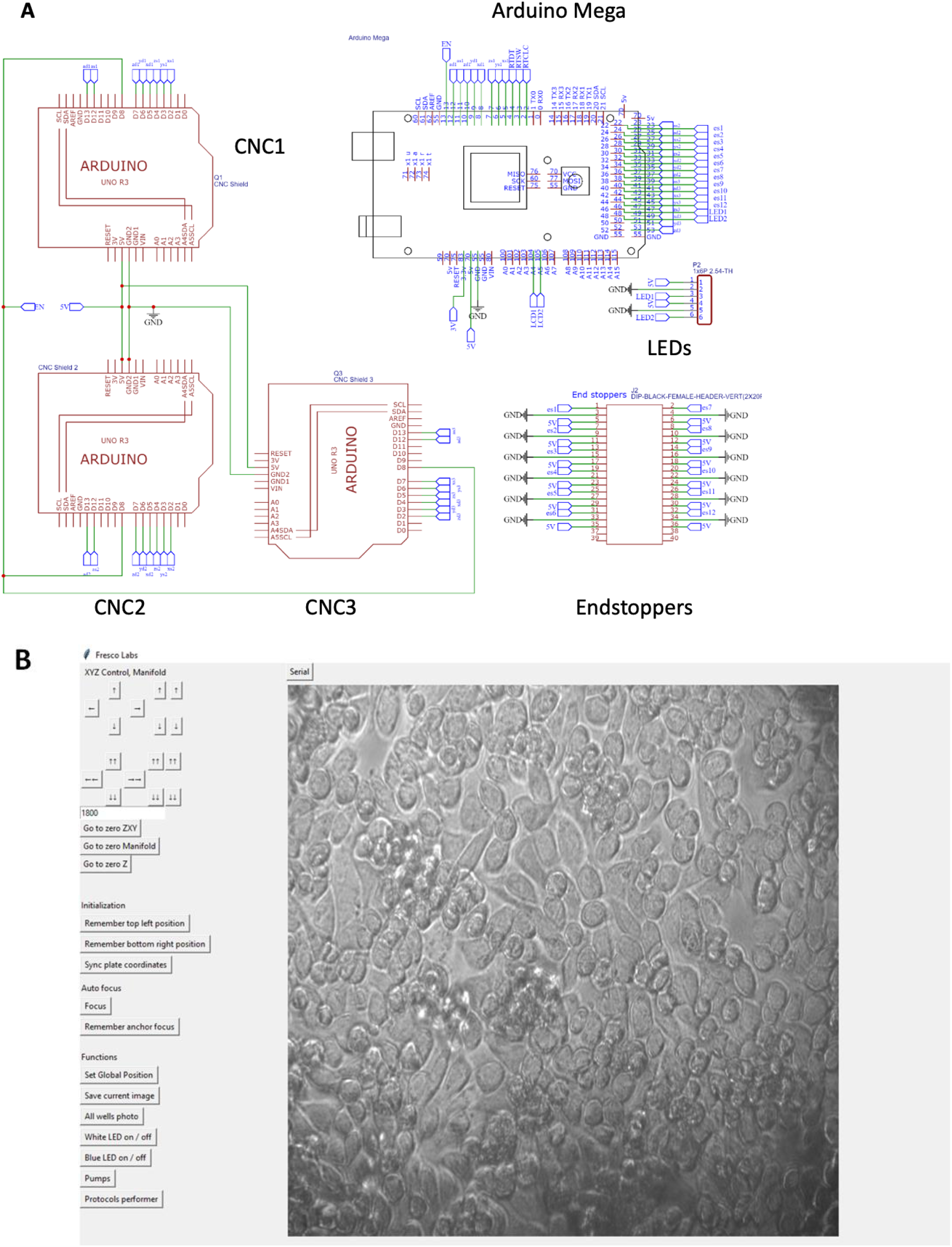
Electric circuits and PCB board controlling the main hardware. A - Schematics of the electric circuit. The main circuit consists of slots for three CNC Shields, Arduino Mega, 12 inputs for end switches. 3 CNC shields (left) drive 12 Nema 17 motors. All 12 motor drivers are set to drive 1/8th step. All three CNC shields are connected to Arduino Mega (top) multiple pins. Bottom – 24 pins are used for 8 endstoppers (3 pins each). The full scheme also containing 12 Volt input from the power source to drive all three CNC shields and LEDs as well as pins reserved for driving LCD screen, SD card and a rotary encoder is shown in Supplementary Figure 3. B – Hardware operating software. Top buttons operate movement of the multiwell plate, perfusion manifold and the focusing system. Only the left side with the buttons is shown. Middle set of buttons drive movement of all motors to zero position. Bottom buttons operate the camera, white and blue LEDs, pumps and run selected protocols.

The microscope is operated by two sets of LEDs – transmitting white light for bright field microscopy and excitation blue (488 nm) light for fluorescent microscopy. Four white LEDs connected in series are then connected to Mosfet IRF520, which in turn is connected to a 12V power supply, ground and pin 46 of the Arduino. The blue LED is operated using a TTL pulse applied to Thorlabs LED driver connected to pin 48 of the Arduino.

Additional pins are reserved for microSD card (Arduino pins 46-52, even numbers), rotary encoder (Arduino pins 2-4) and small LCD display connected to Arduino via i2c protocol (Arduino pins A4, A5). The circuit can be modified to accommodate more CNC shields to increase the overall number of pumps to 22.

#### 1.3 Software

The hardware is operated via an Arduino board that receives instructions from a computer via serial port. Functionality where commands are sent via wifi module or stored in a file in a microSD card are reserved for future versions (all commands are shown in Supplementary Table 1).

The main operating software (Fig. 2B) is written in Python 3 and can be downloaded from https://github.com/frescolabs/FrescoM/blob/master/software. The software is very basic, easy to use and modifiable to fit individual needs. The key functions are as follows:

1. Choose COM port to connect to Arduino (button “Serial”).
2. Move platform forward, backward, left and right. Two sets of buttons allow for large steps or single step movements to be made. The step size can be set.
3. Move application manifold and objective up and down.
4. Return to zero position ╌returns XY platform, the application manifold and Z-focus into the start position.
5. Set top-right and set bottom-right positions of the multiwell plate.
6. Switch ON and OFF white and blue LEDs.
7. Move pumps forwards and backwards. To be used to fill pumps with solutions and cleaning the system after the experiment.
8. Run experiment. Each protocol is implemented as a Python class inherited from a BaseProtocol class (see supplementary text for more information) and overriding the self.perform() function. The following classes should be used for running the key functions:
  a. FrescoXYZ moves multiwell plate in X and Y, manifold, objective and syringe pumps.
  b. ZCamera operates the key camera functions.
  c. ImageStorage saves files generated by the camera.

A few examples of protocol classes are shown in Supplementary Files 1-3.

### 2. System and microscope performance

#### 2.1 Optical resolution

To define the microscope resolution, we used the USAF1951 test chart (Fig. 3A). For 20x objective and under white illumination we can observe the 4th set of elements in group 7 giving us a resolution of at least 181 line pairs per mm. The size of one pixel was 0.26 μm for a 20x objective. These numbers give only approximate values as the platform employs non-infinity corrected objectives and during focusing the distance between the objective and the camera lens may vary.

**Fig 3.**
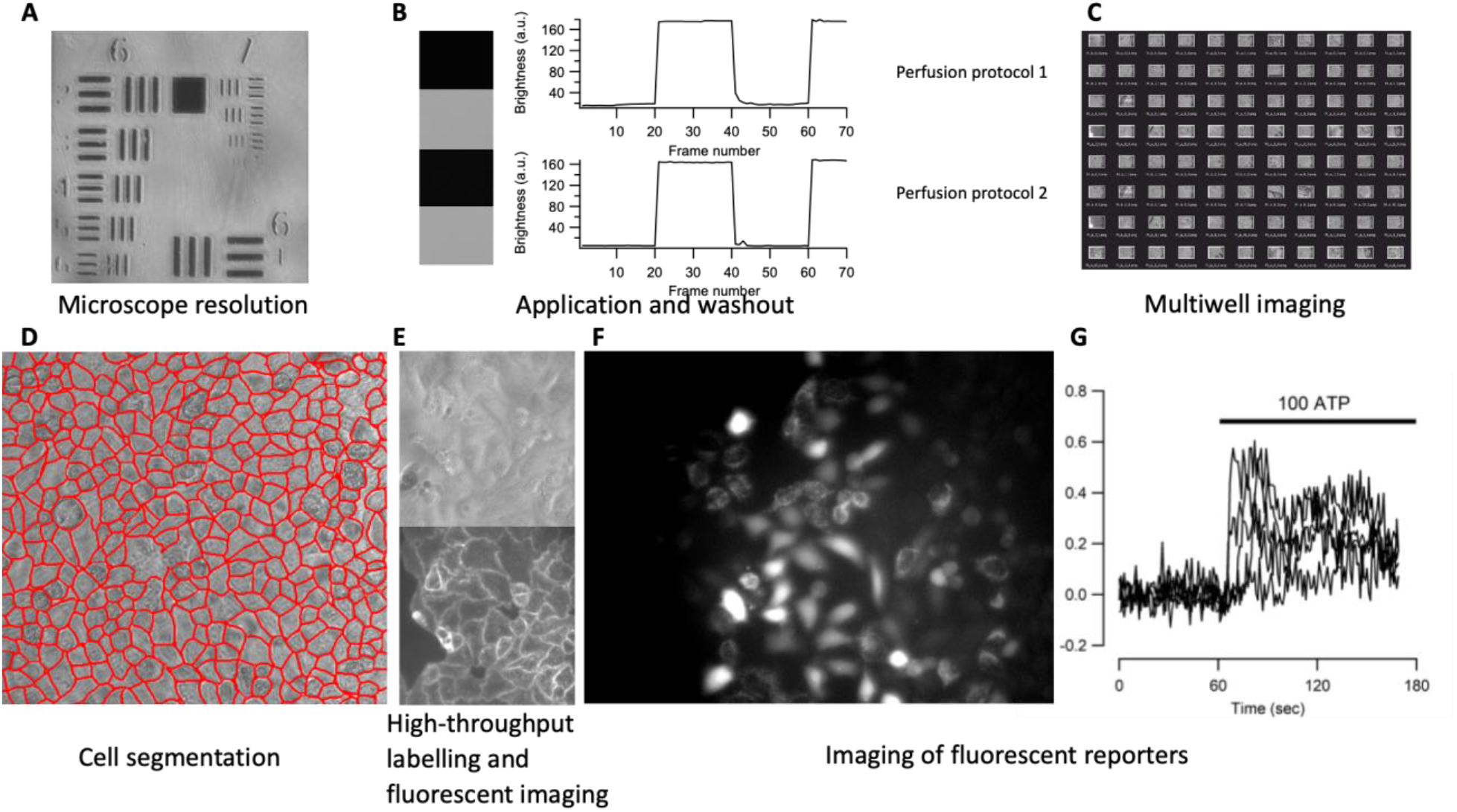
Performance and applicability. A-Micrograph of a USAF 1951 standard. B – Application and washout of fluorescent solution (Lucifer Yellow) recorded by the fluorescent microscope. See details in the main text. Left – frames captured by camera in non-fluorescent and fluorescent solutions. Perfusion Protocol 1 (top) and Perfusion protocol 2 (bottom) exhibit similar degrees of washout. C – example of large scale imaging from a 96 well plate. Cells were fixed in a 96 well plate and imaged under bright light. D – Example of cell segmentation. The software integrates the Cellpose segmentation algorithm [28] and can easily integrate other segmentation algorithms. E – high throughput fluorescent labelling of cells. HeLa cells were automatically labelled with Wheat Germ Agglutinin and washed out with PBS. Top – bright field image. Bottom – fluorescent image of the same cells. F,G - Calcium imaging of HeLa and HCT116 cells. Cells were labelled for 30 min with 1 μM Fluo4-AM (F) or Oregon Green (G) and imaged with sampling frequency of 1 frame per second. F – Example of cells labelled with Fluo4, average of 20 frames. G-5 examples of calcium dynamics in HCT116 cells in response to application of 100 μM ATP. Because the distance between the objective and the camera lens varies from well to well, the scale bar is not shown in any of the experiments.

#### 2.2 System vibration and robustness

To test whether there is any significant vibration in the system we took 50 images one image every 1 second without making the system perform any other tasks (Supplementary protocol 1). We then found the key points in each image using the SIFT algorithm (Supplementary Fig.1) and calculated transformation between the key points using the RANSAC algorithm for each pair of photos (2500 data points altogether). Based on this information we calculated the translation vector and found that the average difference between individual frames was 1.8 pixels and the maximum difference was 6 pixels.

To test how well the system moves from well to well we repeated the same analysis but in this case we moved the multiwell plate from A1 position to H8 and back (Supplementary protocol 2). The plate always went to the same well and approximately the same field of view however the position slightly varied from trial to trial. On average, the offset was 295 pixels, which corresponds to 20% of the field of view. Thus, parallel imaging of multiple well plates requires bringing the plate position to exactly the same position using additional image analysis and programming and/or image registration after the experiment is completed.

#### 2.3 Autofocusing

The designed autofocusing system allows for automatic focusing when moving from well to well. It is based on measuring the value of sharpness for a range of frames, each at a different focal plane. The subsequent positioning of the objective is such that maximum sharpness is achieved. We have implemented and tested 4 different algorithms described in [27] and found that the Tenengrad algorithm calculating the gradient magnitude from a Sobel operator applied to the image to be the most robust for both wide field and fluorescent images. The result of autofocusing performed on different samples is shown on Supplementary Fig.2.

#### 2.4 Perfusion protocols

How well the perfusion system (Fig.1E) exchanges liquids in a well was tested using two different solutions. One solution contained deionised water, while the other contained 25 μg/ml Lucifer Yellow dissolved in deionized water. These two solutions were sequentially applied while taking images by the fluorescent microscope under blue light illumination.

We used two perfusion protocols. In one protocol (Perfusion Protocol 1) we applied the fluorescent solution while sucking the excess using a peristaltic pump connected to a tip located a few millimeters above the other tips. The result of this experiment is shown in Fig. 3B, top. In the second test we used Perfusion protocol 2, in which we positioned one of the applying tips lower than the others, sucked the solution via this tip using a syringe pump and then applied another solution (Fig. 3B, bottom). Both methods yielded a similar degree of washout with Perfusion protocol 2 using smaller volumes but providing less control over the liquid level. Both ways of perfusion have their advantages and can be used in different experimental paradigms. For instance, when one needs quick application of agonist using functional imaging, Perfusion Protocol 1 may be preferable as it offers better control over liquid level. On the other hand, methods such as immunolabelling, particularly using expensive antibodies or other chemicals, may benefit from Perfusion Protocol 2.

### 3. System usability

#### 3.1 High-throughput generation of microscopic images and segmentation

One important advantage of the developed system is that it allows for a large scale generation of images in a large number of wells. To demonstrate this usability, we have generated a Python protocol class (Supplementary file 3) that makes the platform move over all 96 wells, perform focusing on each cell and take a micrograph using either bright field or fluorescent microscopy. Example of such an experiment is shown in Fig. 3C.The micrographs can also be segmented (Fig.3D) using a deep neural network [28] or other algorithms for cell segmentation can be implemented with a minimum of coding.

#### 3.2 Automated labelling

To demonstrate the usability of the developed experimental platform in labelling experiments we used fluorescent Wheat Germ Agglutinin (WGA) that highlights cell membranes. Five rows of wells (40 wells altogether) were automatically washed with PBS and then loaded with a solution containing 5 mkg/mL of fluorescent WGA. After 10 min at room temperature the WGA was washed out with PBS and subjected to fluorescent microscopy. The resulting fluorescent image as well as the corresponding bright field image are shown in Figure 3E. The automatic labelling produces clear images of cells with well-defined plasma membranes thus showing that routing labelling procedures can be automated using the developed platform.

#### 3.3 Imaging fluorescent reporters

Recent advances in fluorescent imaging reporters have allowed researchers to directly monitor signalling pathways in live cells[29]. These include fluorescent reporters for calcium, ERK, NFkB, expression of a number of genes using fluorescent proteins under control of specific promoters and many others. Typical experimental protocol involving fluorescent reporters requires imaging cells for some time before and after application of agonists, growth factors or other active compounds. To demonstrate the usability of the developed experimental platform for such an experimental paradigm, we performed calcium imaging using synthetic fluorescent indicators of calcium concentration. Cells were labelled with fluorescent calcium dye Fluo4-AM (Fig. 3F) or Oregon Green BAPTA-1 AM (Fig. 3G) and subjected to calcium imaging in response to 100 μM ATP (Fig.3G). The quality of the resulting images was sufficient to identify individual cells and calcium response in individual cells can be defined.

## DISCUSSION

There is a high demand for designing affordable and flexible tools for high-scale generation of biological data. Here we report a combination of hardware and software that allows for up to several hundred cell biology experiments performed simultaneously and automatically. Below we discuss applicability of the developed experimental platform and possible ways for its future improvement.

### Importance of automation of biological experiments

Automation of biological experiments is important for two main reasons. First, a large number of data points are required when using modern methods of analysis, such as machine learning and deep learning. A good convolutional neural network algorithm typically requires in the region of 10,000 to 100,000 data points [10]. Considering that there are only a few thousands of cells in a field of view, the same experiments need to be reproduced 10 to 100 times. This number increases significantly if the experimental goal is to optimise conditions for biological experiments.

Second, there is a growing discussion on reproducibility of biological data [12-15]. This is particularly crucial when the results have direct translational applications and can affect future expensive clinical trials. The reproducibility can be improved when the experiments are standardised and when experimentation and data analysis are performed automatically to avoid human errors. The reproducibility of experiments can be further improved when performed on different sources (different cell types, cells with different genetic backgrounds, etc.) further highlighting the necessity of automation of biological experiments.

### Applicability of the developed experimental platform

#### 1. High-throughput screening experiments

Expansion of the compound libraries [16-18] provided a valuable tool to search for new chemicals affecting biological functions. For example, drugs affecting physiological signalling pathways (e.g. GPCRs, etc.) can be assessed by calcium imaging [19-21] or any other forms of functional imaging with a single cell resolution. However, this requires a large number of experiments which is laborious. The framework developed here, easily allows for automatic labelling of cells with fluorescent labels and reporters (Fig.3D-F), application of different agonists (Fig.3B, G) and measurements of calcium signalling (Fig.3G) in a large number of wells.

#### 2. Optimisation experiments

Many biological experiments require optimization of treatment conditions. For example, differentiation of stem cells requires treatment with a large number of growth factors and morphogens at different times and with different dynamics [22, 23]. The framework developed here allows for experiments such as this to be performed in an automatic manner. The outcome of an experiment can then be automatically tested using one of two methods: labeling of cells with synthetic dyes (Fig. 3E), antibodies against appropriate surface markers or a functional experiment (e.g. neurons can be detected by calcium imaging and application of potassium chloride or neurotransmitters).

#### 3. Large scale characterisation of cells derived from individual patients

Substitution of standard cell lines with cells recently derived from individual patients is now becoming increasingly important[24, 25]. The experimental framework developed here would allow a large number of cell lines derived from individual experiments to be tested in a standardized way.

#### 4. Routine cell biology procedures

The developed experimental platform will also be useful in routine lab procedures such as concentration dependence curves, cell dilutions, cytotoxicity studies and others.

#### 5. Data collection for deep learning model training

Tasks such as cell segmentation and classification may be solved by deep learning but they require a large number of cells to be automatically recorded and labelled. The developed platform can be used to image thousands of cells and combine fluorescent and bright field imaging to perform automatic labeling of cell borders and nuclei.

### Further improvements

The modular structure of the platform developed here allows for fairly easy future adjustments to fit individual experimental needs. For example,

1. Substitution of the microscope (Fig.1) with a heat-block will allow to incorporate PCR capability into the analysis. Other means of detection of the experimental outcome (e.g. mini spectrophotometer) will increase the amount experiments that can be performed.
2. Currently, the microscope only provides simple wide-field one colour fluorescence. It can be improved by adding some optical sectioning functionality such as HiLo or light sheet microscopy [26]. The price of the microscope can also be significantly reduced if a custom built LED system is used alongside filter cubes from old fluorescent microscopes. This will reduce the price by £250 and £450 respectively.
3. The number of applied solutions can be increased by redesigning the solution application manifold and the PCB.
4. Cleaning the multi-syringe system can be automated by adding storages for distilled water, alcohol and waste.
5. Experiments can take a long time to perform and cells may need occasional passaging. An algorithm of detecting confluence and then splitting cells into new wells would allow the outcome of experiments to not be affected by overconfluence. In addition to that, selection of individual cells and transferring them into a new well will allow for generation selection of new clones for generation of new lines using transfection or CRISPR.

## MATERIALS AND METHODS

### Cell cultures

HCT116 and HeLa cells were grown in standard Dulbecco modified Eagle’s media (DMEM) supplemented with 10% fetal bovine serum. Cells were passaged once a week when they reached 90% confluence.

### Fluorescent labelling

Fluo4, AM and ORegon Green BAPTA-1 AM (Thermo Fisher Scientific) were dissolved in DMSO (50 μg per 50 μl). 10μl of stock solution and 3 μl of pluronic acid was dissolved in 10 ml, added to the cells. Cells were then incubated at 37 degrees for 45 min. Imaging procedure is described in the main text.

### Licensing and Software Availability

The hardware and software published in work is freely available under Apache 2.0 license. All hardware, firmware, software and Python protocols are available on https://github.com/frescolabs/FrescoM

## ACKNOWLEDGEMENTS

Microscopes were designed and built by AN and AC and supported by the University of Sheffield Alumni Fund. The mechanics and electronics was designed and implemented by PK and AN. We thank Elliot Birkett for reading and comments on the manuscript.

## References

1. Hinton, B., Ma, L., Mahmoudzadeh, A.P., Malkov, S., Fan, B., Greenwood, H., Joe, B., Lee, V., Kerlikowske, K., and Shepherd, J. (2019). Deep learning networks find unique mammographic differences in previous negative mammograms between interval and screen-detected cancers: a case-case study. Cancer Imaging 19, 41.

2. Hinton, G. (2018). Deep Learning-A Technology With the Potential to Transform Health Care. JAMA 320, 1101–1102.

3. LeCun, Y., Bengio, Y., and Hinton, G. (2015). Deep learning. Nature 521, 436–444.

4. Hilsenbeck, O., Schwarzfischer, M., Loeffler, D., Dimopoulos, S., Hastreiter, S., Marr, C., Theis, F.J., and Schroeder, T. (2017). fastER: a user-friendly tool for ultrafast and robust cell segmentation in large-scale microscopy. Bioinformatics 33, 2020–2028.

5. Coudray, N., Ocampo, P.S., Sakellaropoulos, T., Narula, N., Snuderl, M., Fenyo, D., Moreira, A.L., Razavian, N., and Tsirigos, A. (2018). Classification and mutation prediction from non-small cell lung cancer histopathology images using deep learning. Nat Med 24, 1559–1567.

6. Van Valen, D.A., Kudo, T., Lane, K.M., Macklin, D.N., Quach, N.T., DeFelice, M.M., Maayan, I., Tanouchi, Y., Ashley, E.A., and Covert, M.W. (2016). Deep Learning Automates the Quantitative Analysis of Individual Cells in Live-Cell Imaging Experiments. PLoS Comput Biol 12, e1005177.

7. Kusumoto, D., and Yuasa, S. (2019). The application of convolutional neural network to stem cell biology. Inflamm Regen 39, 14.

8. Heras, F.J.H., Romero-Ferrero, F., Hinz, R.C., and de Polavieja, G.G. (2019). Deep attention networks reveal the rules of collective motion in zebrafish. PLoS Comput Biol 15, e1007354.

9. https://www.biorxiv.org/content/10.1101/2020.03.20.000133v1.

10. Mahoney Deep Learning vs. Traditional Computer Vision. https://arxiv.org/ftp/arxiv/papers/1910/1910.13796.pdf.

11. Razlivanov, I., Liew, T., Moore, E.W., Al-Kathiri, A., Bartram, T., Kuvshinov, D., and Nikolaev, A. (2018). Long-term imaging of calcium dynamics using genetically encoded calcium indicators and automatic tracking of cultured cells. Biotechniques 65, 37–39.

12. Freedman, L.P., Cockburn, I.M., and Simcoe, T.S. (2015). The Economics of Reproducibility in Preclinical Research. PLoS Biol 13, e1002165.

13. Ioannidis, J.P. (2005). Why most published research findings are false. PLoS Med 2, e124.

14. Ioannidis, J.P. (2007). Why most published research findings are false: author’s reply to Goodman and Greenland. PLoS Med 4, e215.

15. Ioannidis, J.P. (2014). How to make more published research true. PLoS Med 11, e1001747.

16. Cases, M., Garcia-Serna, R., Hettne, K., Weeber, M., van der Lei, J., Boyer, S., and Mestres, J. (2005). Chemical and biological profiling of an annotated compound library directed to the nuclear receptor family. Curr Top Med Chem 5, 763–772.

17. Rickardson, L., Fryknas, M., Haglund, C., Lovborg, H., Nygren, P., Gustafsson, M.G., Isaksson, A., and Larsson, R. (2006). Screening of an annotated compound library for drug activity in a resistant myeloma cell line. Cancer Chemother Pharmacol 58, 749–758.

18. Root, D.E., Flaherty, S.P., Kelley, B.P., and Stockwell, B.R. (2003). Biological mechanism profiling using an annotated compound library. Chem Biol 10, 881–892.

19. Berridge, M.J. (2001). The versatility and complexity of calcium signalling. Novartis Found Symp 239, 52–64; discussion 64-57, 150-159.

20. Berridge, M.J., Bootman, M.D., and Roderick, H.L. (2003). Calcium signalling: dynamics, homeostasis and remodelling. Nat Rev Mol Cell Biol 4, 517–529.

21. Bootman, M.D., Berridge, M.J., and Roderick, H.L. (2002). Calcium signalling: more messengers, more channels, more complexity. Curr Biol 12, R563–565.

22. Kim, J.H., Auerbach, J.M., Rodriguez-Gomez, J.A., Velasco, I., Gavin, D., Lumelsky, N., Lee, S.H., Nguyen, J., Sanchez-Pernaute, R., Bankiewicz, K., et al. (2002). Dopamine neurons derived from embryonic stem cells function in an animal model of Parkinson’s disease. Nature 418, 50–56.

23. Panchision, D.M., and McKay, R.D. (2002). The control of neural stem cells by morphogenic signals. Curr Opin Genet Dev 12, 478–487.

24. Rominiyi, O., Al-Tamimi, Y., and Collis, S.J. (2019). The ‘Ins and Outs’ of Early Preclinical Models for Brain Tumor Research: Are They Valuable and Have We Been Doing It Wrong? Cancers (Basel) 11.

25. Stringer, B.W., Day, B.W., D’Souza, R.C.J., Jamieson, P.R., Ensbey, K.S., Bruce, Z.C., Lim, Y.C., Goasdoue, K., Offenhauser, C., Akgul, S., et al. (2019). A reference collection of patient-derived cell line and xenograft models of proneural, classical and mesenchymal glioblastoma. Sci Rep 9, 4902.

26. https://www.microscopyu.com/techniques/light-sheet/light-sheet-fluorescence-microscopy.

27. Bueno-Ibarra M.A. et al. (2005). Fast autofocus algorithms for automated microscopes. Optical Engineering 44(6), 063601

28. Stringer C. et al. (2021). Cellpose: a generalist algorithm for cellular segmentation. Nat Methods 18:100–106

29. Nasu Y. et al., Structure- and mechanism guided designs of single fluorescent-protein based biosensors. (2021). Nat Chemical Biol https://doi.org/10.1038/s41589-020-00718-

