## Supplementary Text for "An open-source experimental framework for automation of high-throughput cell biology experiments"

This file contains assembly instructions, software user guide and supplementary figures referenced in the main text.

**Assembly instructions**

Building the system requires some time (approximately 2 days), concentration, calmness and basic soldering skills.

MakerbeamXL extrusions are connected to each other using aluminium L- and T-shaped brackets. Each bracket is screwed by 5 M3 screws. Think in advance where nuts have to be placed as forgetting a nut may require disassembly of some parts. Below we provide schematic instruction for building the hardware and the electric circuits. The positions of L- and T- brackets are not shown but they can be put anywhere as long as they properly bind extrusions together. Figures below show how to assemble the platform. Black arrows show where you need to insert the nuts **prior to frame assembly.**

Step 1. 3D print all parts required for the hardware. All STL files can be found at https://github.com/frescolabs/FrescoM/tree/master/hardware/print Generate PCB (the scheme and/or Gerber file can be downloaded from https://github.com/frescolabs/FrescoM/tree/master/hardware/pcb).

Step 2. Main frame and the X-axis.

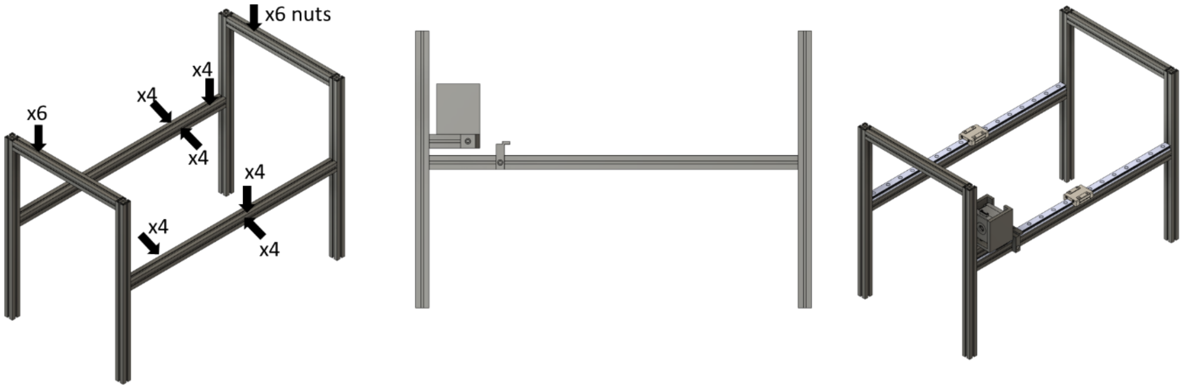


Assemble the main frame, attach X-motor extrusion, X-motor holder and X-axis endstop holder as shown in figures above. Screw the endstop to the holder using M3 screws. Attach MGH12 rails. **NB! Remember to insert M3 nuts to the top of lower horizontal extrusions, at least 4 nuts to each side of the lower horizontal extrusions and 4 screws to the inner side of each upper horizontal extrusions before assembling the frame!**

Step 3. Y-axis assembly

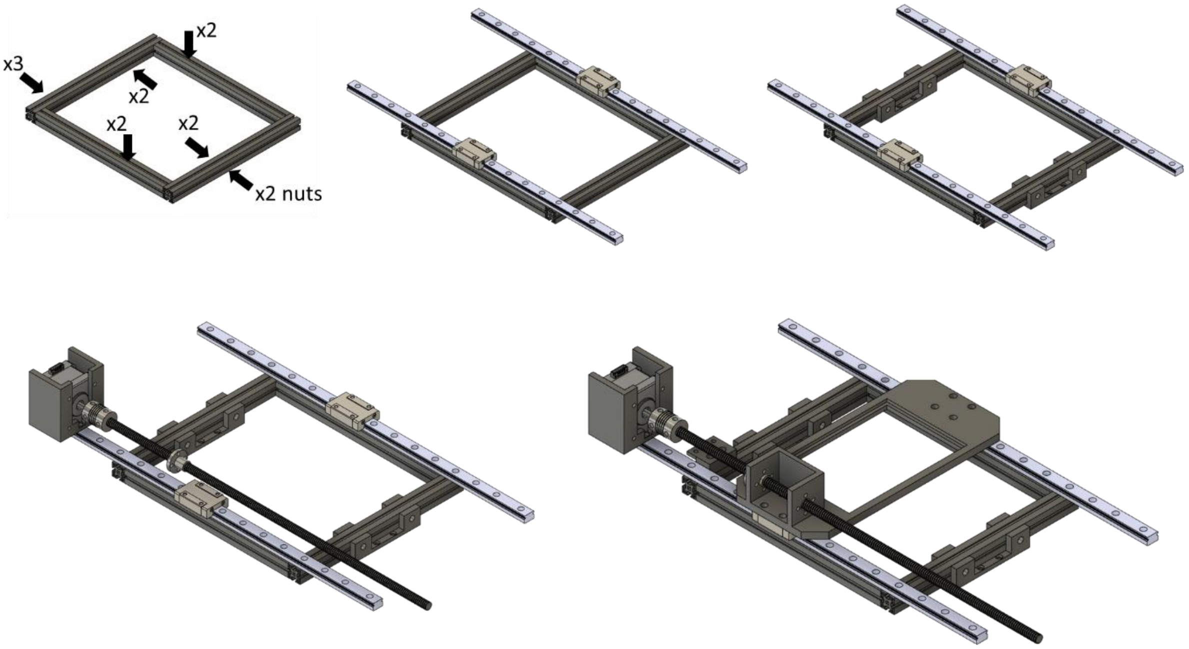


Assemble the Y-axis as shown above. Bind the main frame extrusions using L-shapes positioned below the frame. Attach one endstop to the endstop holder. **NB! Add nuts to each hidden extrusion side.**

Step 4. Perfusion manifold assembly.

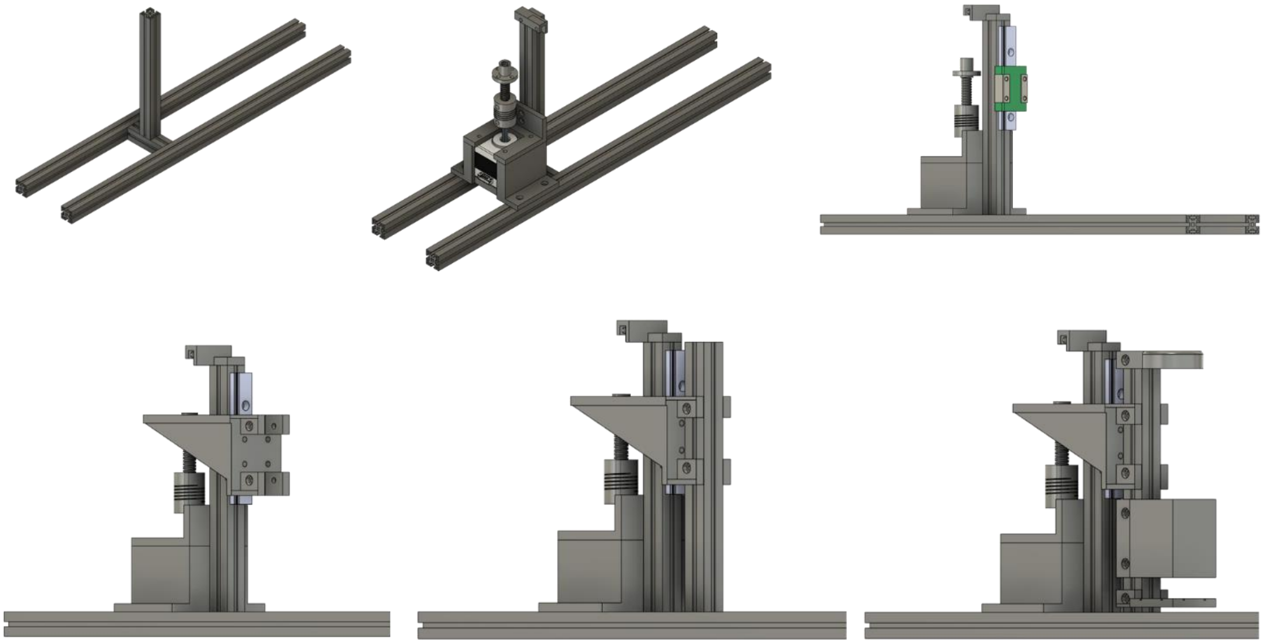

Assemble the perfusion manifold as shown above. Attach the endstop to the endstopholder (top of the vertical extrusion). Before mounting the LED holder at the bottom, insert 4 white LEDs, solder them to be connected in sequence and solder unconnected electrodes to a pair of long wires (see Figure below).

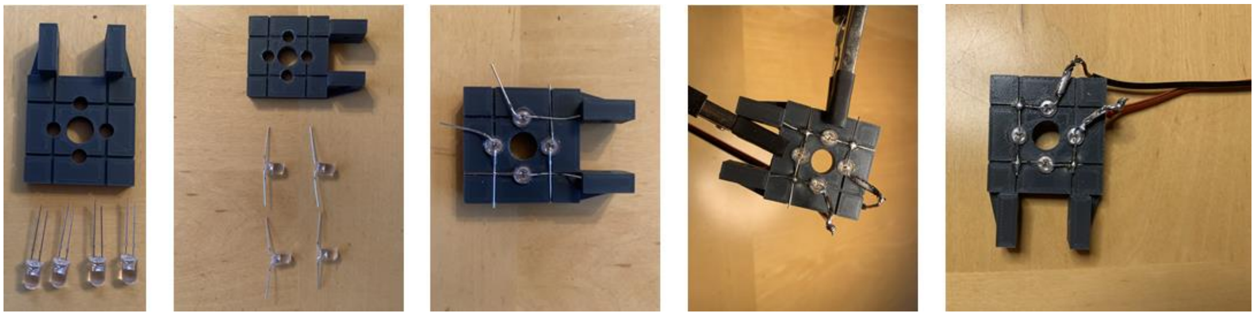

Step 5. Mini microscope assembly

The following instruction describes how to assemble a small fluorescent microscope using Thorlabs details. The cost of the current configuration is £2000. This can be reduced by £450 if a cheaper LED and constant current driver is used. See https://github.com/Frescolabs/FrescoM for additional instructions.

1. Connect one mirror cube with two or 4 assemble rods
2. Connect filter cube
3. Insert 1 dichroic mirror into the filter cube as described in Thorlabs website
4. Insert excitation and emission filters into short SM1 tubes, fix with SM1RR retaining rings and connect the filter cube
5. Insert the 100mm focal distance lens to 100 mm SM1 tube and connect to the short emission filter SM1 tube.
6. Insert one 20 mm lens into a short SM1 tube and connect to the excitation filter to focus excitation light onto the sample
7. Connect mounted LED (e.g. M470L4)
8. Connect the mini-microscope to the Z-focusing system

Step 6. Focusing system assembly

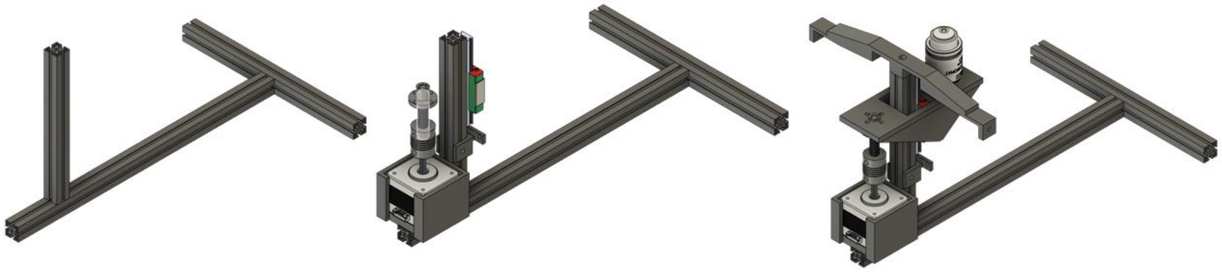

Assemble the focusing system as shown above. Attach endstop to the endstop holder.

Step 7. Assemble the microscope and connect everything together as shown below.

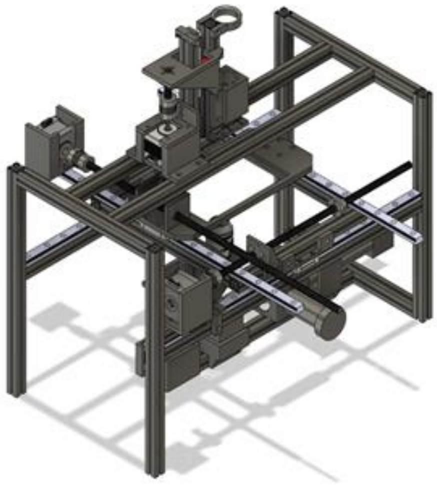

Step 8. Connect syringe pumps and/or peristaltic pumps via tubings and gel loading tips.

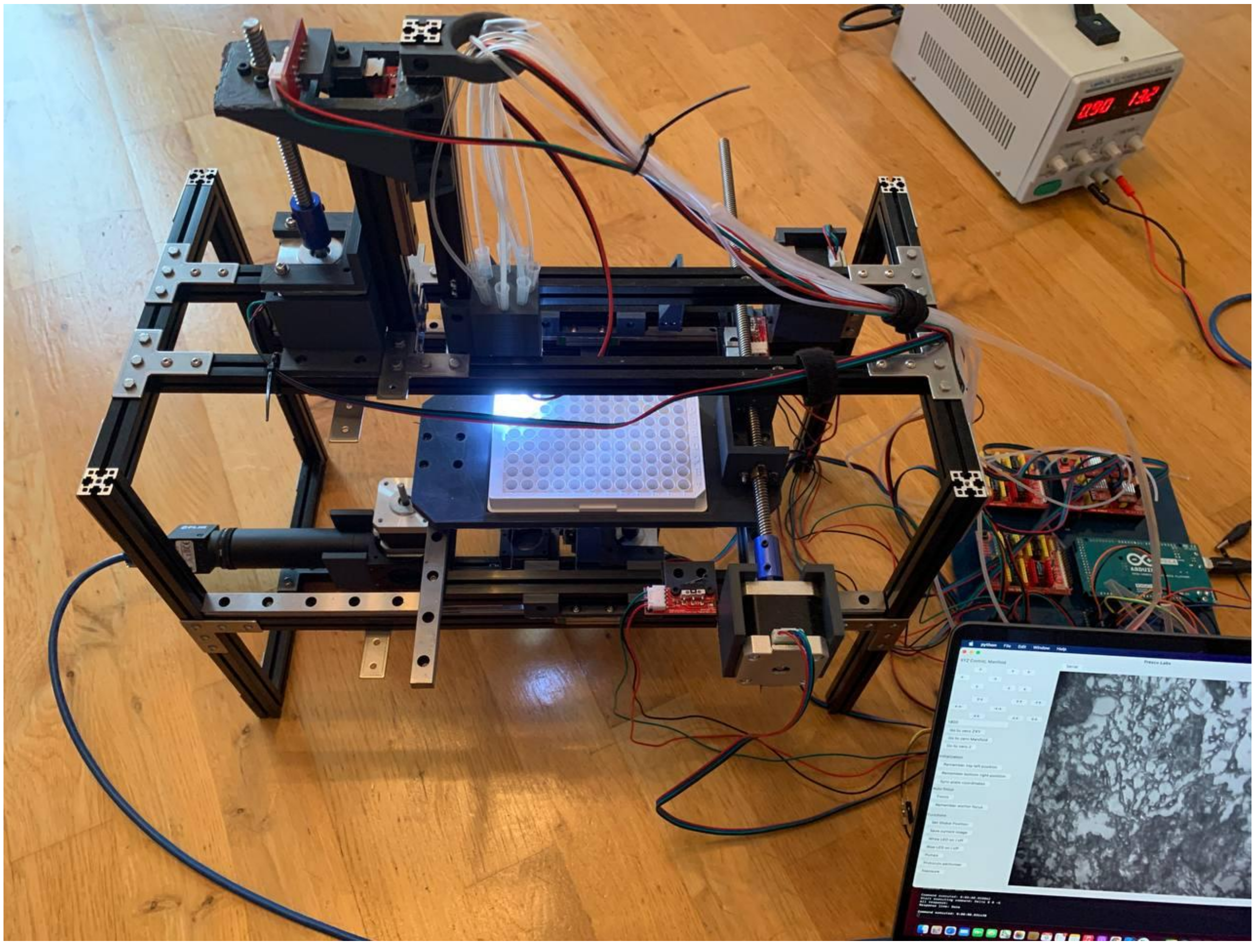

Step 9. Electric circuit assemble

We provide all files that can be used to order PCB. All pins need to be soldered by yourself. However, some companies provide this service if you are not willing to solder parts by yourself.

1. Order PCB from preferred supplier (e.g. EasyEDA)
2. Optional: solder all pin slots to the PCB
3. Insert Arduino Mega and all three CNC shields.
4. Connect CNC shields with jumpers as shown in figure below and then insert A4988 drivers.
5. Connect 12V to each CNC shield and to the PCB as shown on figure above.
6. Connect each endstop as shown on figure above. First connect endstops for X-, Y-, autofocus- and perfusion manifold in this order.
7. Connect white LEDs as shown in figure below
8. Connect blue LED driver as shown in figure above. Connect the other side to Thorlabs LED driver.

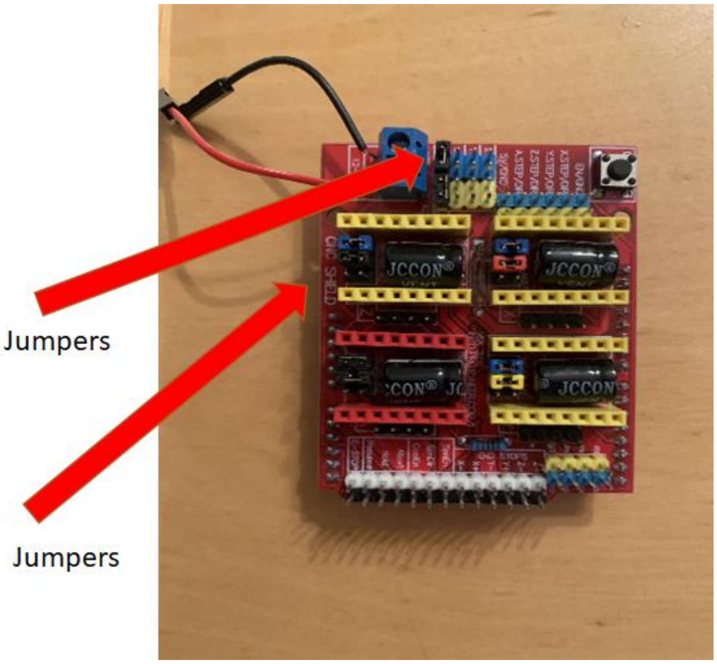


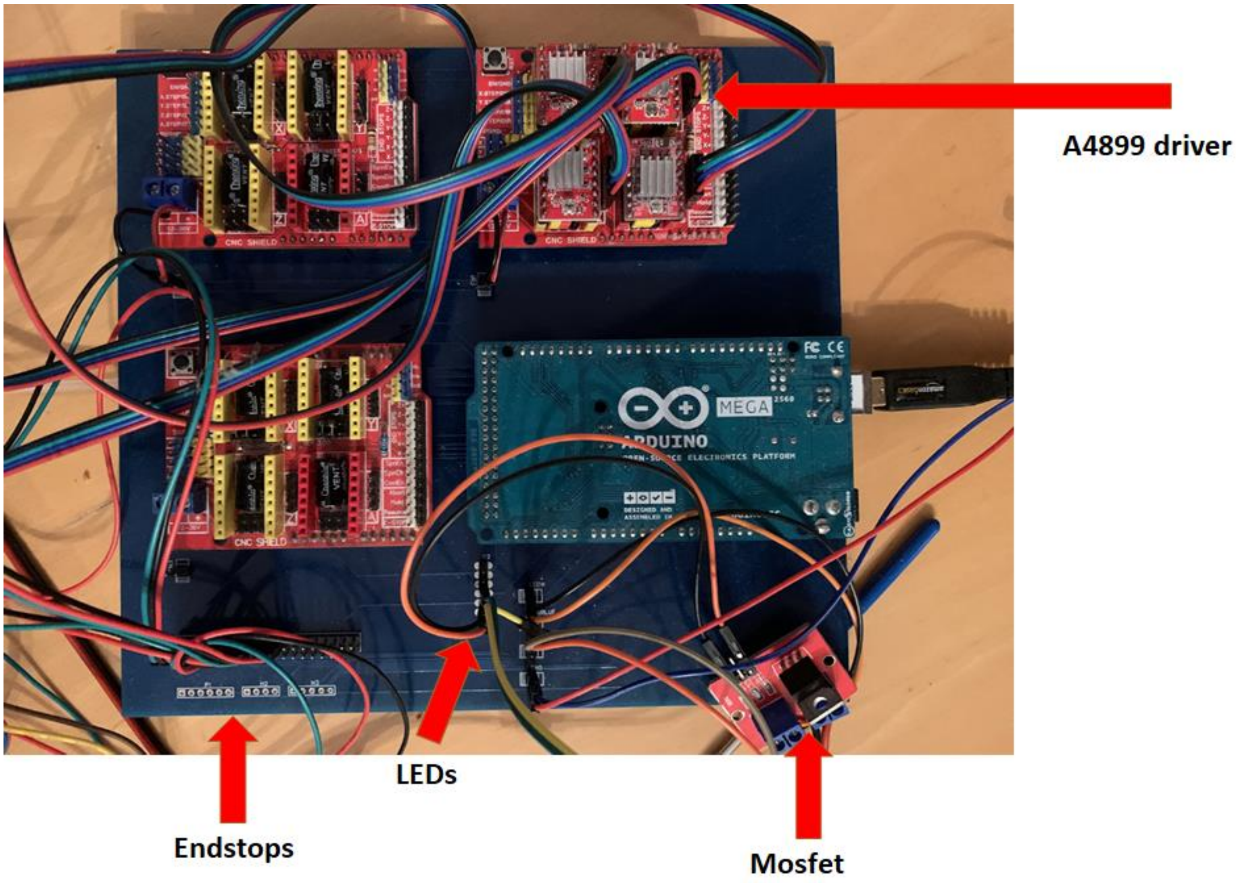


**Firmware**

The hardware is operated via an Arduino board that receives instructions from a computer via serial port. Functionality where commands are sent via wifi module or stored in a file in a microSD card are reserved for future versions.  The commands shown in Table 1.

| **Command** | **Action** |
|---|---|
| Zero | Move X and Y to zero position |
| VerticalZero | Move Z to Zero position |
| Position (e.g. Position 1000 1000 1000) | Set position |
| Delta (e.g. Delta 0 100 0) | Move X, Y, Z by the given number of steps |
| RememberTopLeft | Remember top left |
| RememberBottomRight | Remember bottom right |
| GetTopLeftBottomRightCoordinates | Send coordinates for the plate |
| ManifoldZero | Move Manifold to zero position |
| DeltaPump 0 -100 | Move pump # 0 for -100 steps |
| ManifoldDelta | Moves manifold |
| SwitchLedW | Switch on and off top white LED |
| SwitchLedB | Switch on and off blue LED for microscope |
| GetCurrentPosition | Returns current XYZ |

**Supplementary Table 1. List of commands used by Arduino to operate the main hardware.** The Arduino code can be downloaded from https://github.com/frescolabs/FrescoM/blob/master/firmware

**Protocol programming guide using Python**

The following part describes the main Python classes to be used to generate new protocols. Python has become popular among biologists and therefore, instead of development of complex GUI where new protocols can be set up, we propose programming protocols using Python.

The basics of protocol programming is simple:

1. Generate a new class inherited from BaseProtocol class
2. Override the perform() function
3. Override the constructor
4. Save the new script in software/services/protocols folder. Class name should have the same name as the file but with Snake Case translated to Pascal case. For example, if you name your class AllWellsPhotoProtocol save it in all_wells_photo_protocol.py file.
5. Press “Protocol Performer” button on the main Form, chose the protocol and run it.

Additional steps:
1. Import the following classes:

```
from services.protocols.base_protocol import BaseProtocol
from services.fresco_xyz import FrescoXYZ
from services.z_camera import ZCamera
from services.images_storage import ImagesStorage
```

2. Override the constructor in the following way:

```
def __init__(self,
             fresco_xyz: FrescoXYZ,
             z_camera: ZCamera,
             images_storage: ImagesStorage):
    super(AllWellsPhotoProtocol, self).__init__(fresco_xyz=fresco_xyz,
                                                z_camera=z_camera,
                                                images_storage=images_storage)
    self.images_storage = images_storage
```

where AllWellsPhotoProtocol is the name of your class.

3. Use the following functions
**self.fresco_xyz.white_led_switch()** to switch on and off the white LED
**self.fresco_xyz.go_to_zero_manifold()** to move manifold to zero position
**self.z_camera.focus_on_current_object()** to run autofocus
**self.hold_position()** to wait for some time (in seconds)
**self.z_camera.fresco_camera.get_current_image()** to capture an image
**self.fresco_xyz.manifold_delta()** to move manifold certain number of steps
**self.fresco_xyz.delta (x, y, z)** to move XY stage and the focusing system by X, Y or Z steps. If you used hardware and electronics configuration described above move X or Y by 1800 steps to move to the next well.
**self.fresco_xyz.delta_pump ()** to move the given pump by given number of steps
**self.fresco_xyz.blue_led_switch()** to switch on and off the white LED

### SUPPLEMENTARY FIGURES

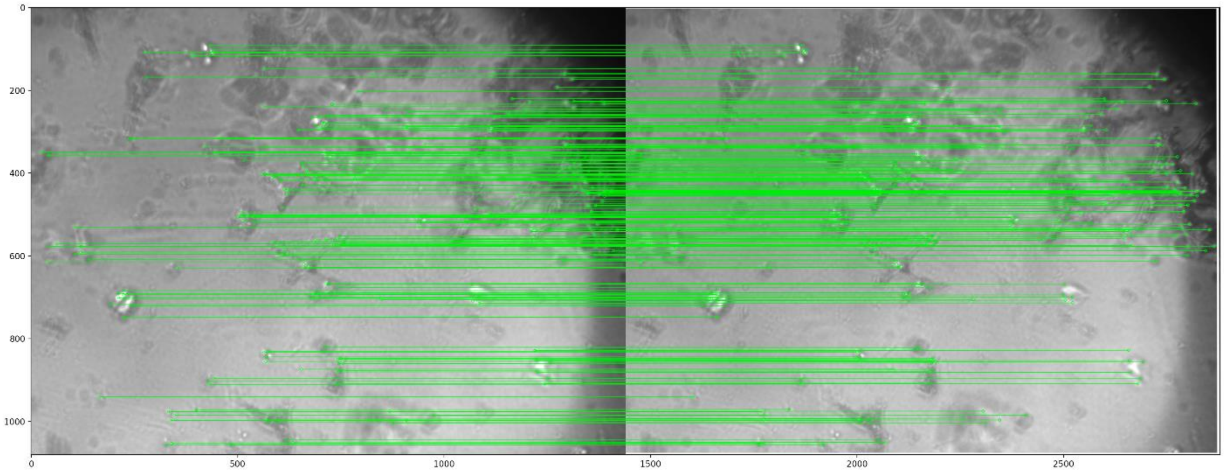


**Supplementary Figure 1. Calculation system vibration using SIFT and RANSAC algorithms.** Two frames were subjected to scale invariant feature transform (SIFT) that detects the key points of each image (green circles). Then the RANSAC algorithm was used to compare the position of each point.

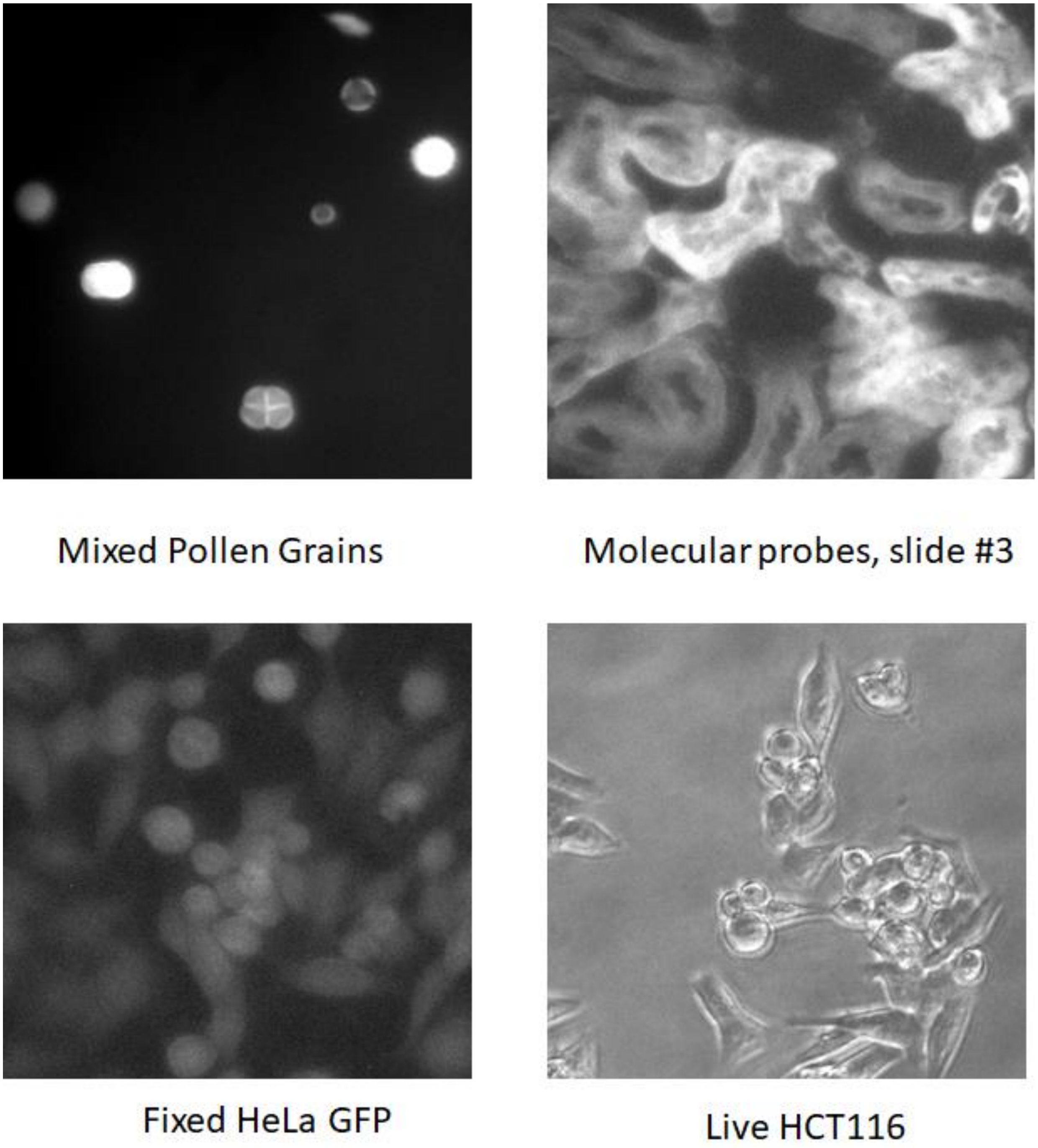


**Supplementary Figure 2. Examples of autofocusing performed on different types of images.**

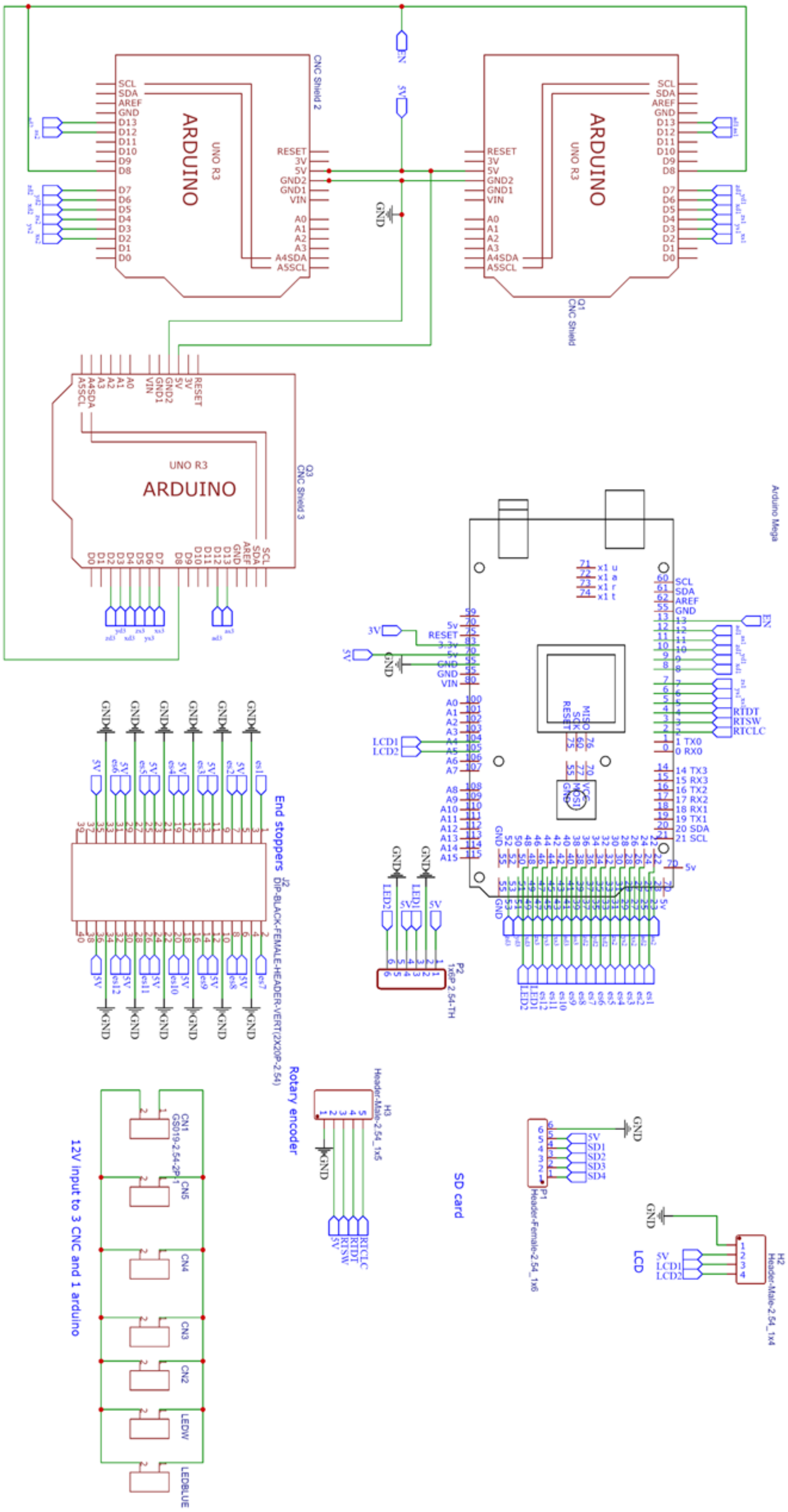


**Supplementary Figure 3. Full size electronic scheme**
